## Supplementary information for "A generalizable brain extraction net (BEN) for multimodal MRI data from rodents, nonhuman primates, and humans"

**Supplementary Materials List**

Fig. S1. Performance comparison of BEN with two benchmark settings on the task of cross-modality domain transfer.
Fig. S2. Performance comparison of BEN with two benchmark settings on the task of cross-platform domain transfer.
Fig. S3. BEN outperforms traditional SOTA methods in functional MRI scans acquired from multiple species on different magnetic field strengths.
Fig. S4. Linear regression coefficients and Bland–Altman analysis demonstration of the rest datasets.
Fig. S5. Error maps of BEN and SOTA methods.
Fig. S6. Execution time comparison of BEN with other methods.
Fig. S7. BEN's uncertainty is consistent with interrater' disagreement.
Fig. S8. Interrater disagreement.
Fig. S9. BEN provides a measure of uncertainty that potentially reflects the disagreement of conventional toolboxes in human data.
Fig. S10. BEN's transferability is not dependent on specific source dataset.
Fig. S11. UMAP visualizes BEN's transfer learning.

Table S1. MRI scan information of the fifteen animal datasets and three human datasets.
Table S2. Performance comparison of BEN with SOTA methods on the source domain (Mouse-T2WI-11.7T).
Table S3. Performance comparison of BEN with SOTA methods on two public datasets.
Table S4. Ablation study of each module of BEN in the source domain.
Table S5. Ablation study of each module of BEN in the target domain.
Table S6. BEN provides interfaces for the following conventional neuroimaging software.
Table S7. Protocols and parameters used for conventional neuroimaging toolboxes.

### Supplementary Figures

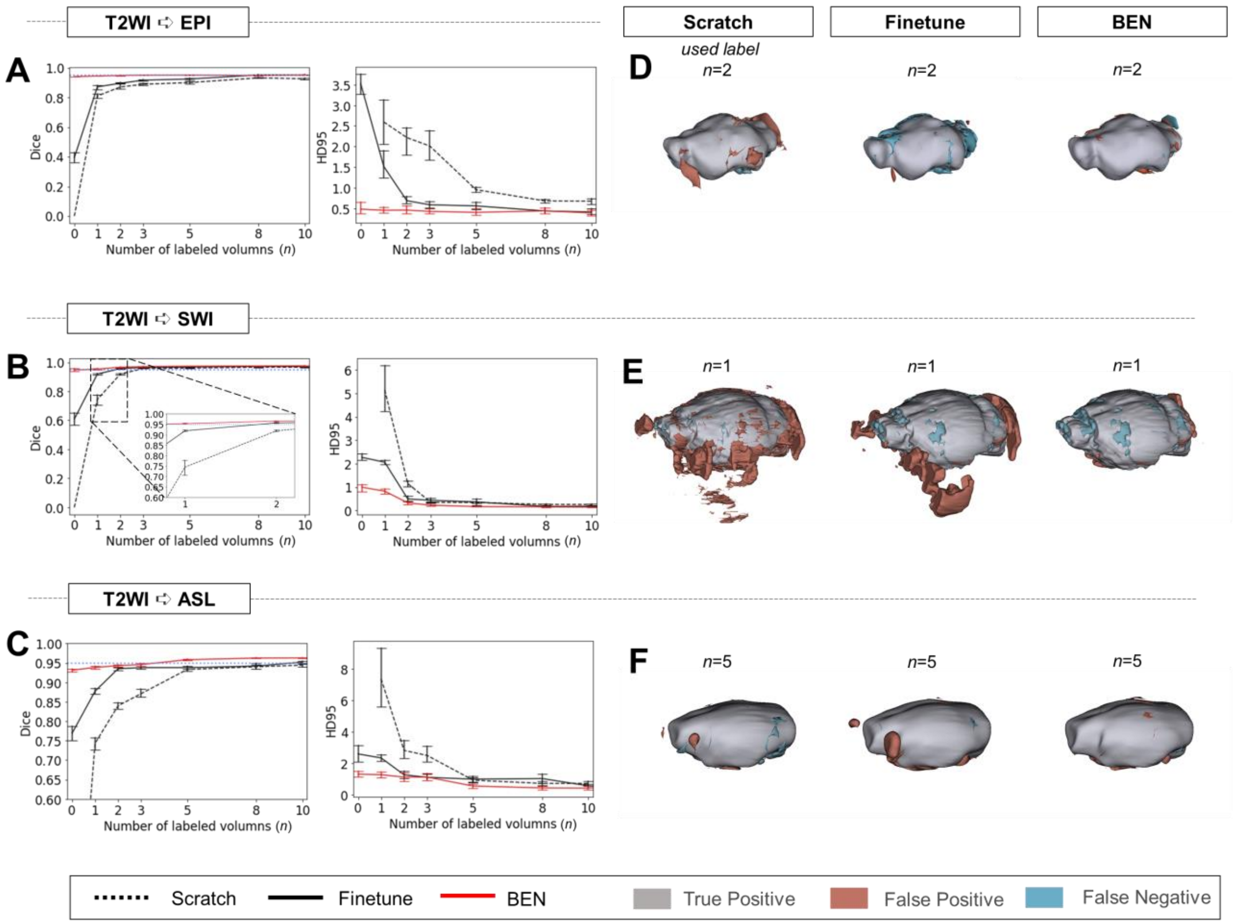


**Figure S1. Performance comparison of BEN with two benchmark settings on the task of cross-modality domain transfer.**

**(A - C)** The curve plots representing domain transfer tasks across modality, showing the variation in the segmentation performance in terms of the Dice score and 95% Hausdorff distance (HD95) (y axis) as a function of the number of labeled volumes (x axis) for training from scratch (black dotted lines), fine-tuning (black solid lines) and BEN (red lines). From top to bottom: **(A)** T2WI to EPI, **(B)** T2WI to SWI and **(C)** T2WI to ASL with an increasing amount of labeled training data (*n*=1, 2, 3, 5, 8, 10, as indicated on the x axis in each panel). BEN consistently surpasses other methods with less labeled data required. Note that, BEN achieves acceptable performance via zero-shot inference (*n*=0), where no label is used in the target domain. In this case, fine-tuning directly infer on target datasets and training from scratch failed to perform. Error bars represent the mean with a 95% confidence interval (CI) for all curves. The blue dotted line corresponds to *y(Dice)*=0.95, which can be considered to represent qualified performance for brain extraction. **(D - F)** 3D renderings of representative segmentation results (the number *(n)* of labels used for each method is indicated in each panel). Images with fewer colored regions represent better segmentation results. (gray: true positive; brown: false positive; blue: false negative.)

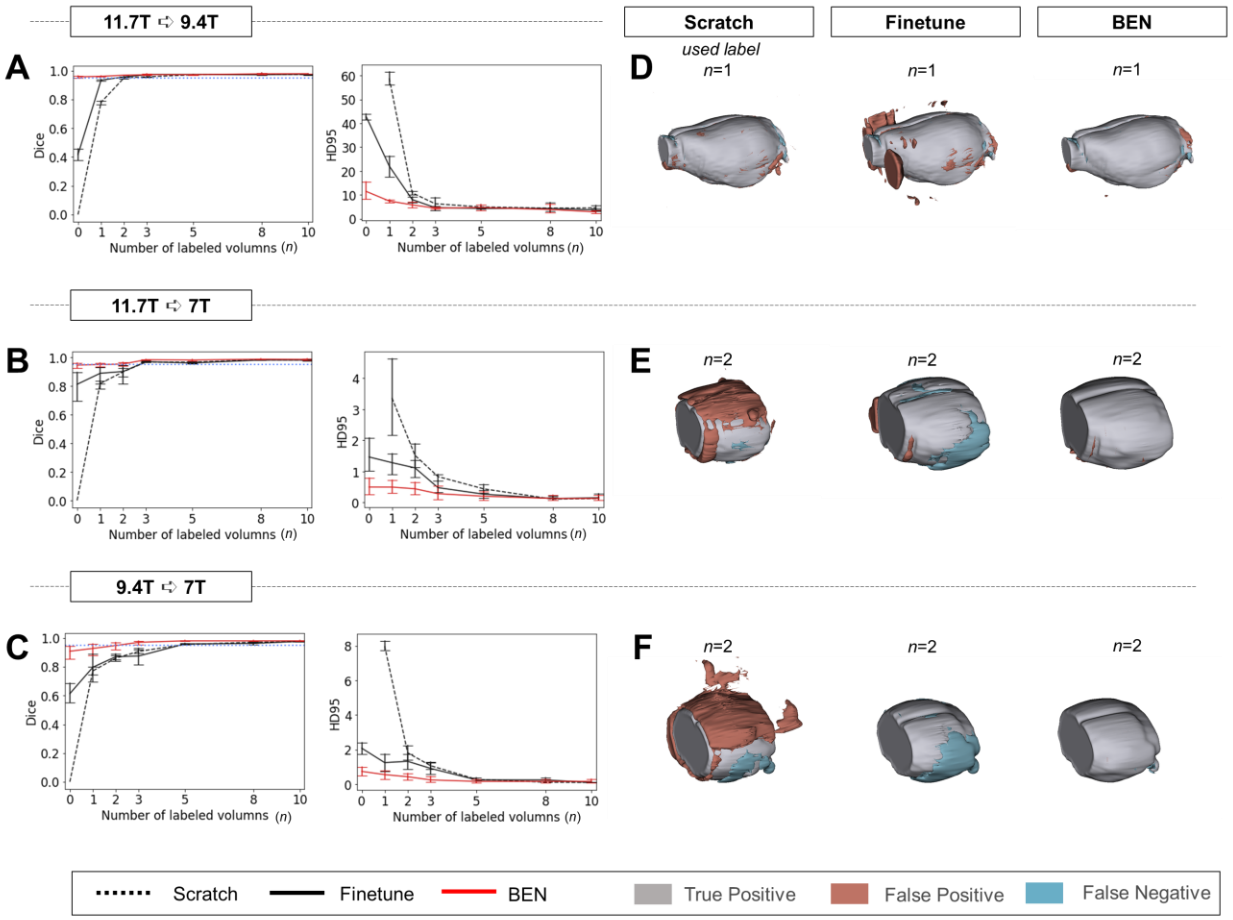


**Figure S2. Performance comparison of BEN with two benchmark settings on the task of cross-platform domain transfer.**

**(A - C)** Curve plots representing domain transfer tasks across platforms with different magnetic field strengths, showing the variation in the segmentation performance in terms of the Dice score and 95% Hausdorff distance (HD95) (y axis) as a function of the number of labeled volumes (x axis) for training from scratch (black dotted lines), fine-tuning (black solid lines) and BEN (red lines). From top to bottom: **(A)** 11.7T to 9.4T, **(B)** 11.7T to 7T and **(C)** 9.4T to 7T with an increasing amount of labeled training data (*n*=1, 2, 3,..., as indicated on the x axis in each panel). BEN reaches saturation point much earlier in all three tasks than other methods, indicating that fewer labels are required for BEN than for other methods. Error bars represent the mean with a 95% confidence interval (CI) for all curves. The blue dotted line corresponds to *y(Dice)*=0.95, which can be considered to represent qualified performance for brain extraction. **(D - F)** 3D renderings of representative segmentation results (the number *(n)* of labels used for each method is indicated in each panel). Note that due to MRI acquisition, the anterior and posterior of brains in the 7T dataset are not included. Images with fewer colored regions represent better segmentation results. (gray: true positive; brown: false positive; blue: false negative.)

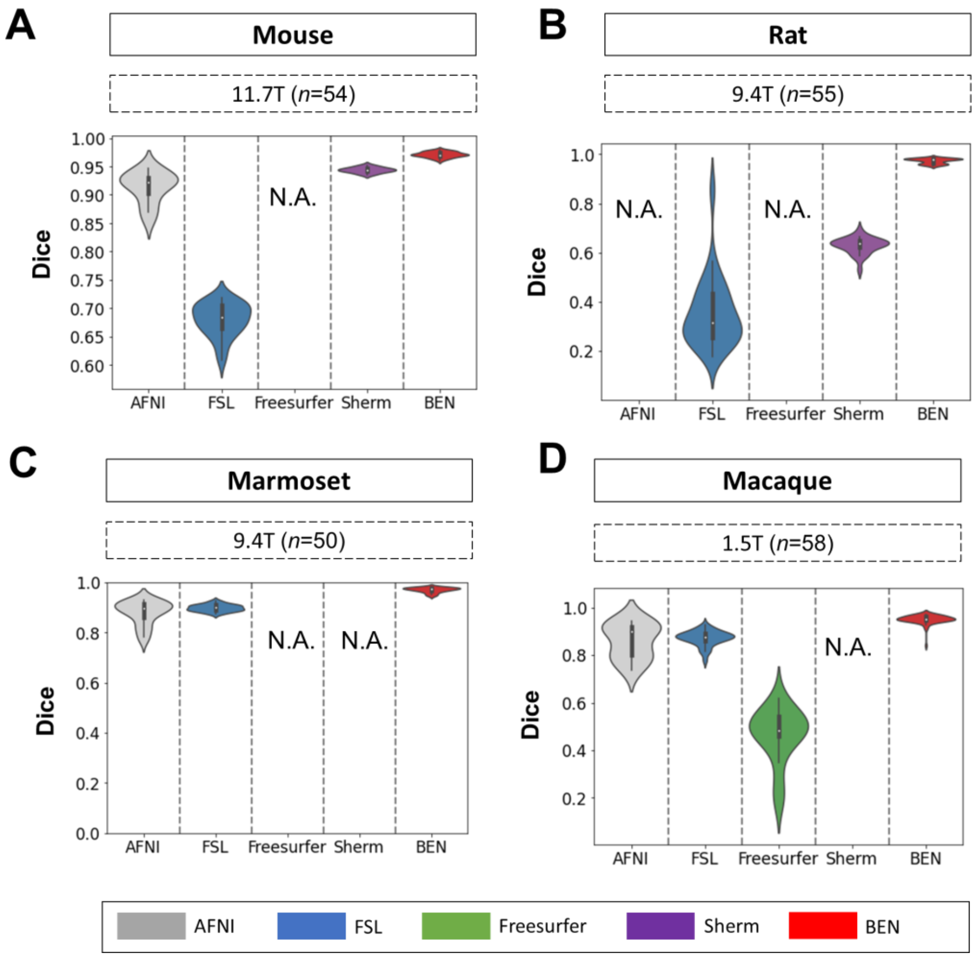


**Figure S3. BEN outperforms traditional SOTA methods in functional MRI scans acquired from multiple species on different magnetic field strengths.**

**(A - D)** Violin plots and inner box plots showing the Dice scores of each method for **(A)** mouse (*n*=54), **(B)** rat (*n*=55), **(C)** marmoset (*n*=50), and (d) macaque (*n*=58) MRI images acquired. The field strength is illustrated with different markers above each panel, and the results for each method are shown in similar hues. The median values of the data are represented by the white hollow dots in the violin plots, the first and third quartiles are represented by the black boxes, and the interquartile range beyond 1.5 times the first and the third quartiles is represented by the black lines. 'N.A.' indicates failure of the method on the corresponding dataset. (gray: AFNI; blue: FSL; green: Freesurfer; purple: Sherm; red: BEN).

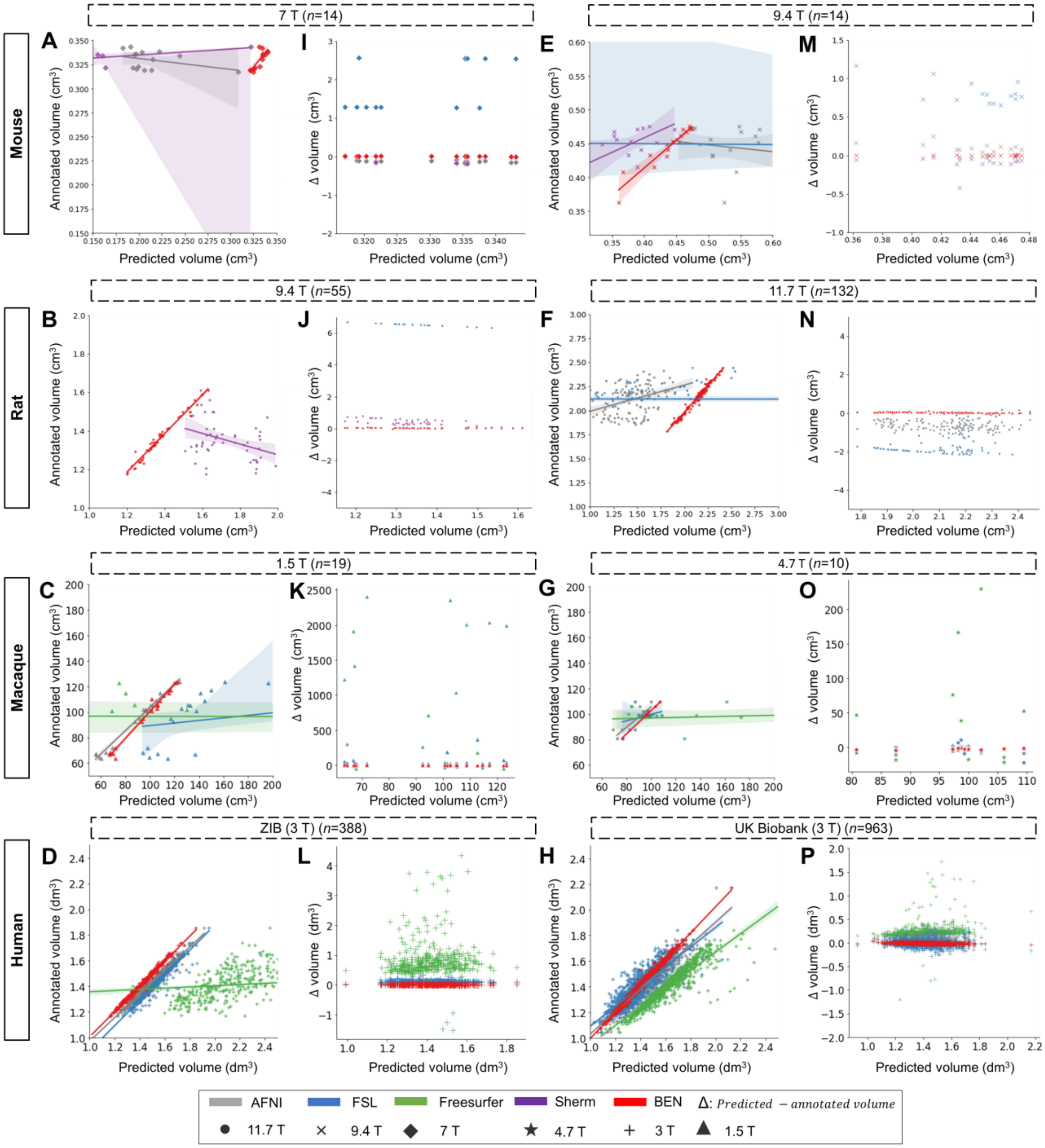

**Figure S4. Linear regression coefficients and Bland–Altman analysis demonstration of the rest datasets.**

**(A-D, E-H)** The segmentation accuracy is assessed using the linear regression coefficients (LRCs) (*n* of volumes used are shown above the figure panels). Compared with other methods, BEN achieves better performance. The magnetic field strengths are also shown in different markers according to the legend. Each dot in a graph represents one sample in the dataset. The error bands represent 95% CI in plots. **(I - L, M - P)** Bland–Altman analysis showing high consistency between the BEN results and expert annotations. The biases between two observers and the 95% CIs for the differences are shown in the tables above each plot. Each method is shown using the same hue in all plots (gray: AFNI; blue: FSL; green: FreeSurfer;

purple: Sherm; red: BEN). Different field strengths are shown with different markers (●: 11.7 T; ×: 9.4 T; ◆: 7 T; ★: 4.7 T; +: 3 T; ▲: 1.5 T).

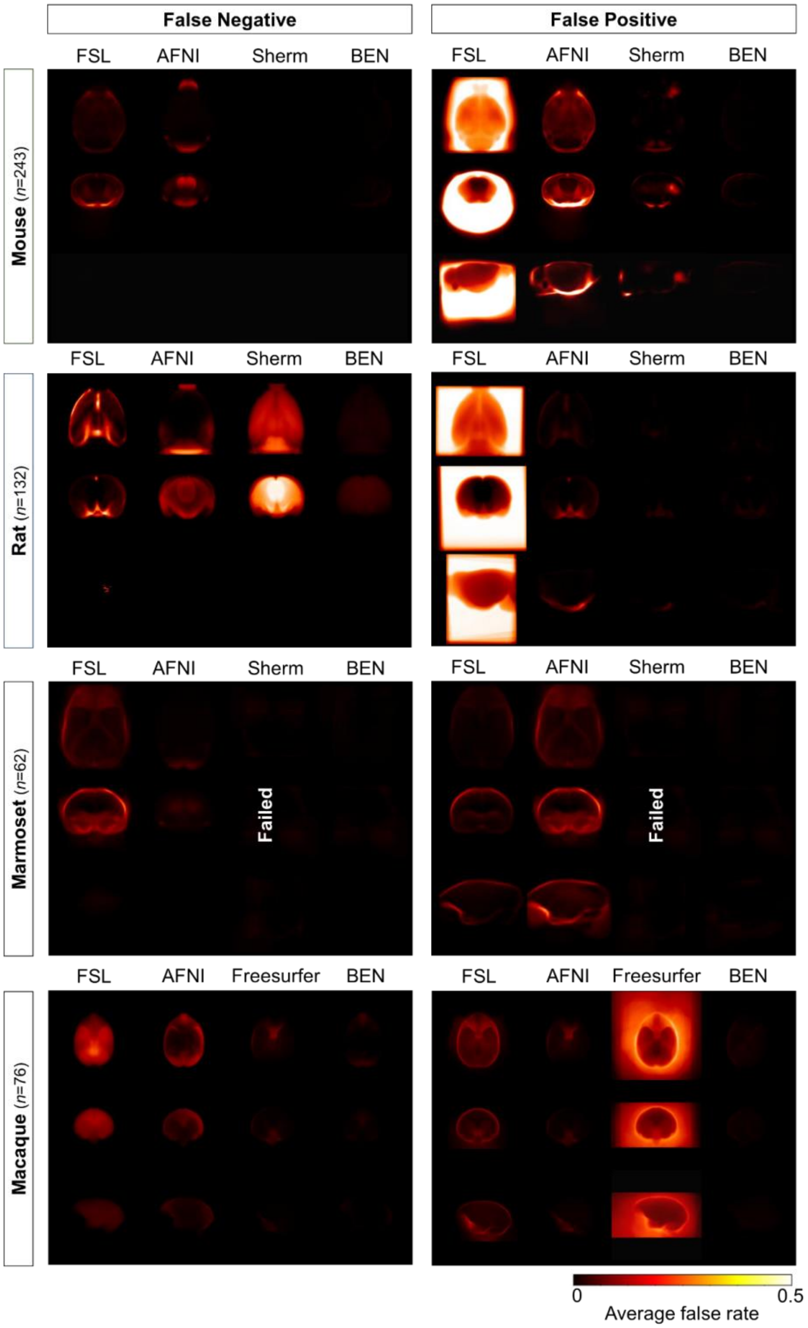


**Figure S5. Error maps of BEN and SOTA methods.**
Heatmap projections of average false negative (FN) and false positive (FP) for each species (mouse, rat, marmoset, and macaque). From the first row to the third row: axial, coronal, and sagittal view. Compared with other methods, BEN shows much less FN and FP errors. The upper extreme represents a high systematic number of FNs and FPs. (mouse: *n*=243 in mouse-T2WI-11.7T dataset; rat: *n*=132 in rat-T2WI-11.7T dataset; marmoset: *n*=62 in marmoset-T2WI-9.4T dataset; macaque: *n*=76 in macaque-T1WI dataset).

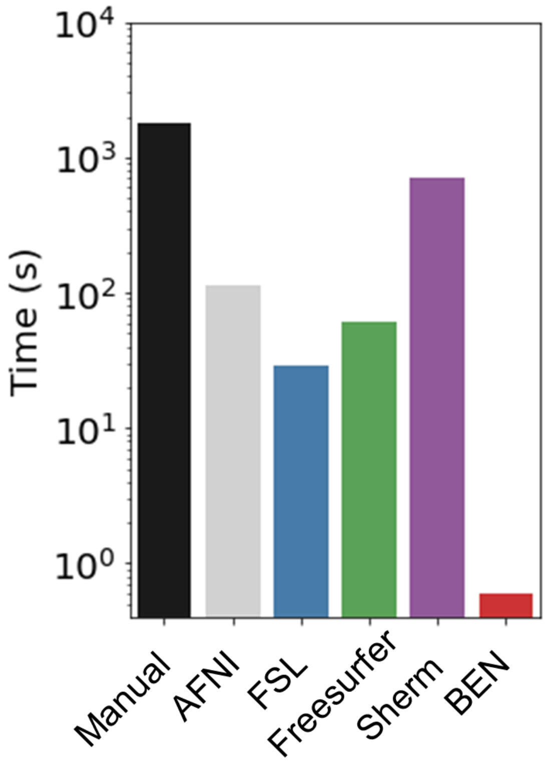


**Figure S6. Execution time comparison of BEN with other methods.**
Compared with other conventional methods, BEN has unequaled faster processing speed. The plot is in log scale and represents average processing time to segment one 3D volume.

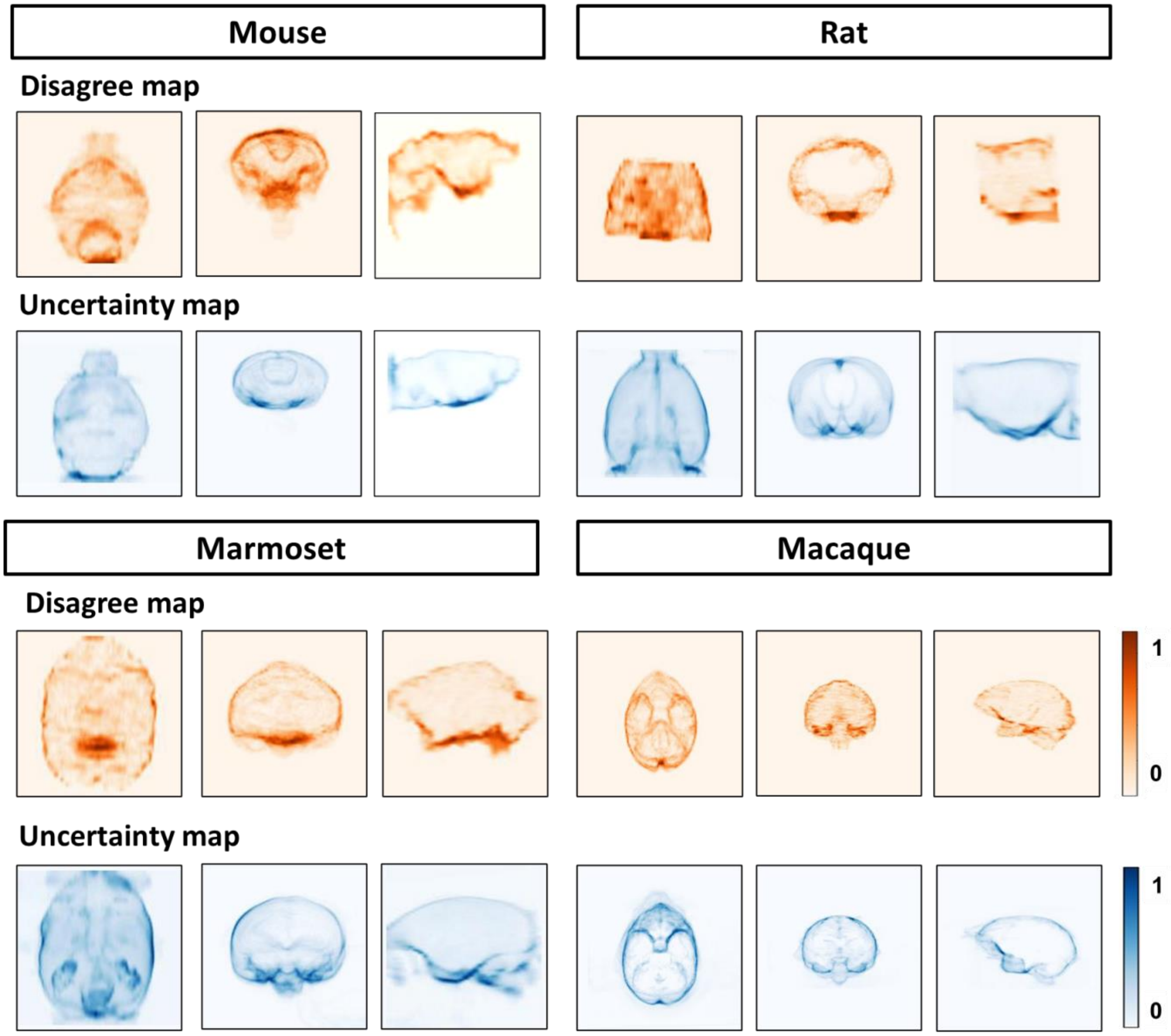


**Figure S7. BEN's uncertainty is consistent with interrater' disagreement.**
Heat map projections of raters' disagreement (orange hues) and BEN uncertainty (blue hues) across four species using the ground truth as the reference. From left to right: Sagittal, coronal and axial view. These disagree maps and uncertainty maps share similar feature distribution patterns.

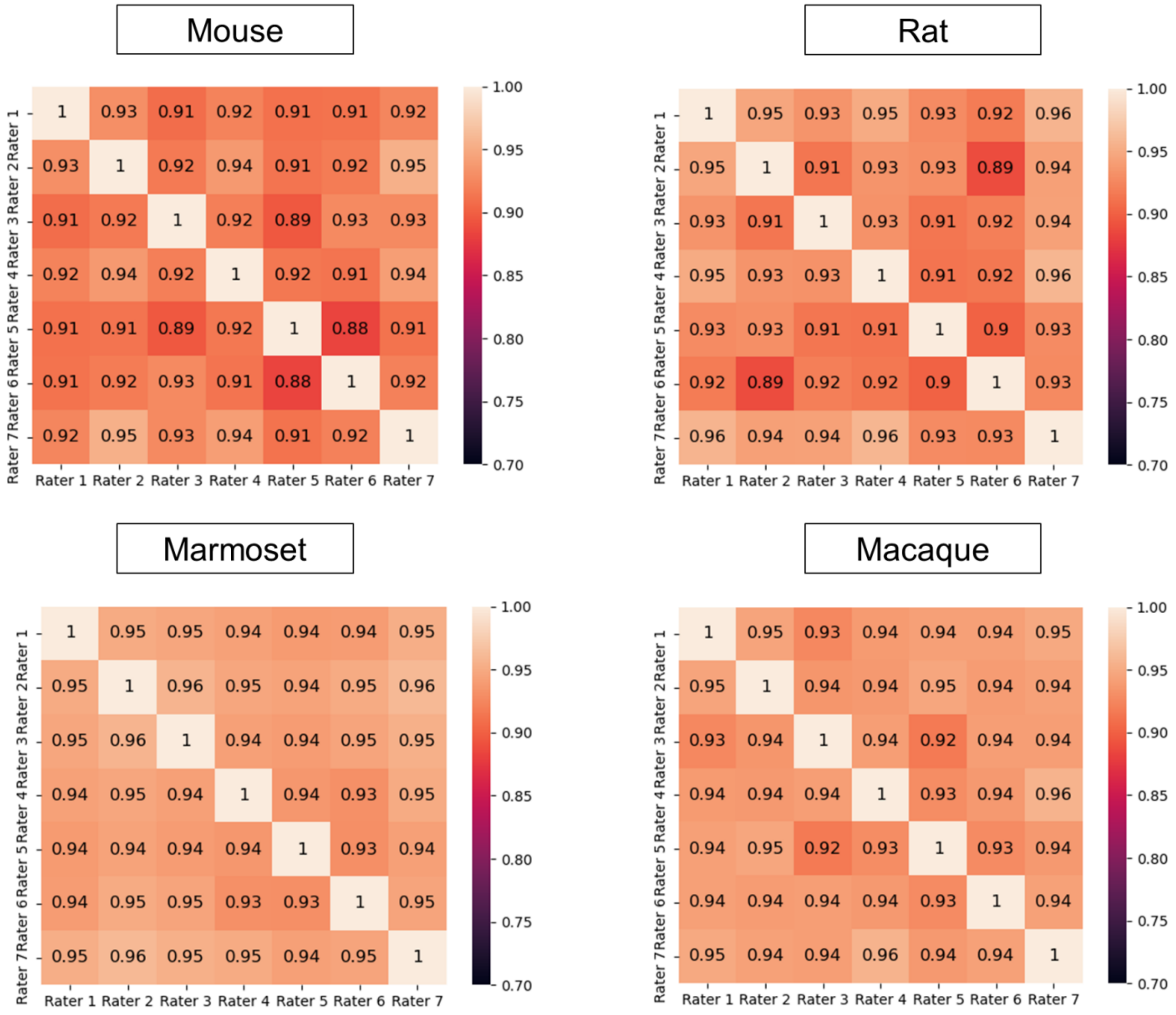


**Figure S8. Interrater disagreement.**
Heatmaps representing the segmentation Dice scores of each junior rater as calculated using the labels from one of the others as the ground truth on the rat images and the macaque images.

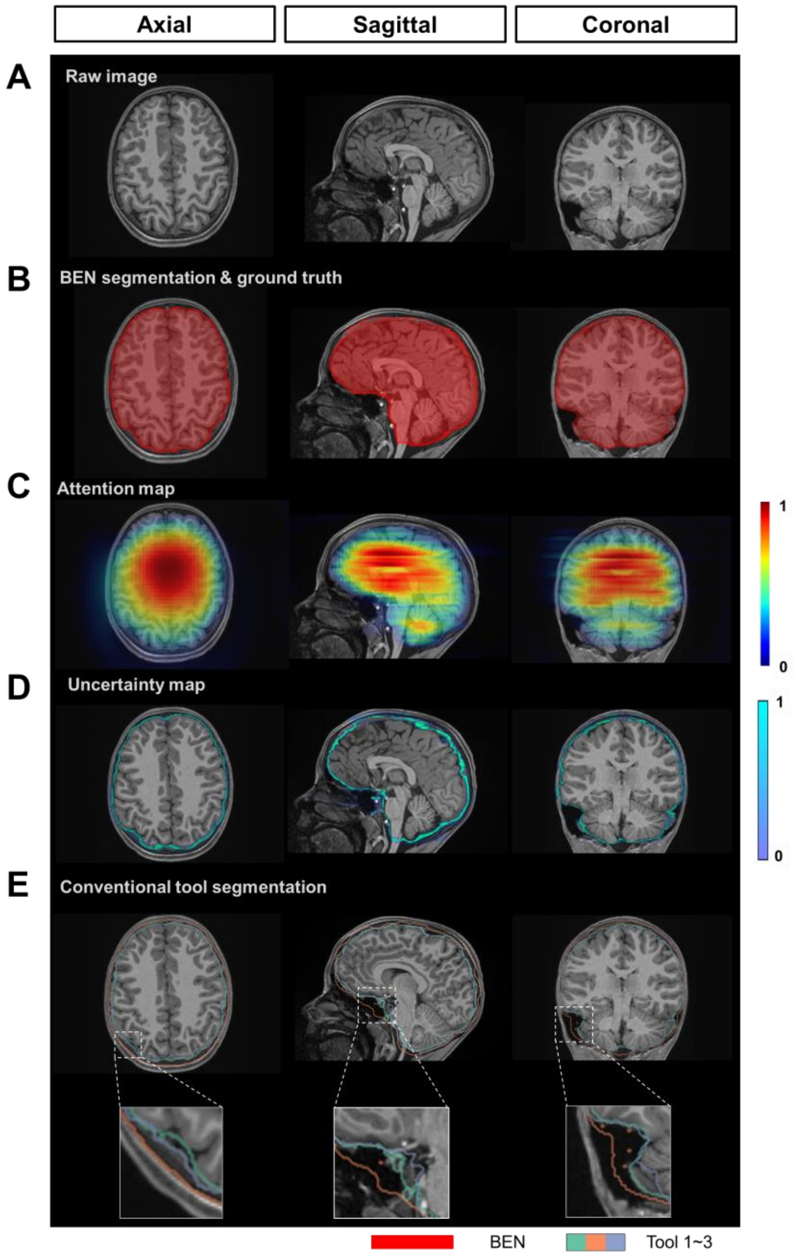


**Figure S9. BEN provides a measure of uncertainty that potentially reflects the disagreement of conventional toolboxes in human data.**
**(A)** Representative structural MR images of the human in the ABCD dataset (*n*=3,250). **(B)** BEN's segmentation (red semi-transparent areas) overlaid with annotation of experts' consensus, which is considered as ground truth (red opaque lines). **(C)** Attention map shows the key semantic features in images where BEN captures. **(D)** Uncertainty map shows the regions where BEN has less confidence. The uncertainty values are normalized (0 ~ 1) for better visualization. **(E)** Segmentations obtained by toolboxes (*n*=3) are shown in different color opaque lines.

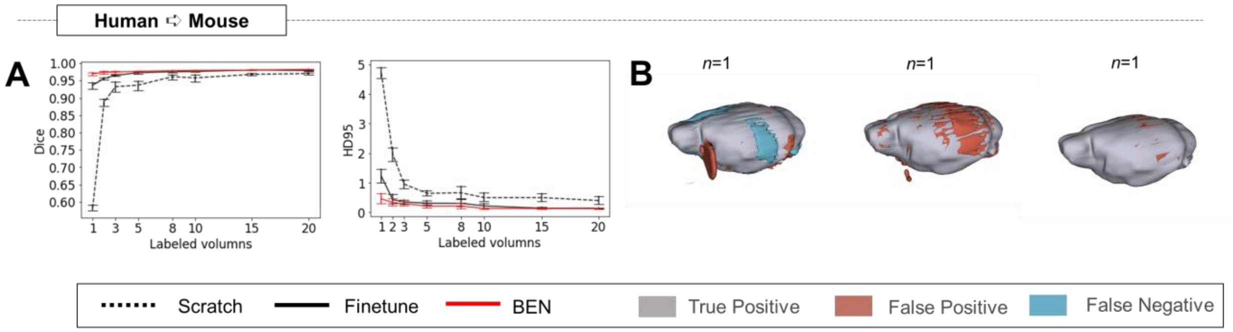


**Figure S10. BEN's transferability is not dependent on specific source dataset.**
In our previous experiments, the mouse-T2WI-11.7T dataset was used as the source domain and transferred the model to other species, including humans. Here, we reselect the human as the source domain and transfer the model to the mouse dataset. **(A)** Human to Mouse with an increasing amount of labeled training data (*n*=1, 2, 3,..., as indicated on the x axis in each panel). Curve plots representing domain transfer tasks across species, showing the variation in the segmentation performance in terms of HD95 (y axis) as a function of the number of labeled volumes (x axis) for training from scratch (black dotted lines), fine-tuning (black solid lines) and BEN (red lines). **(B)** 3D renderings of representative segmentation results (the number *(n)* of labels used for each method is indicated in each panel). Images with fewer colored regions represent better segmentation results. (gray: true positive; brown: false positive; blue: false negative.)

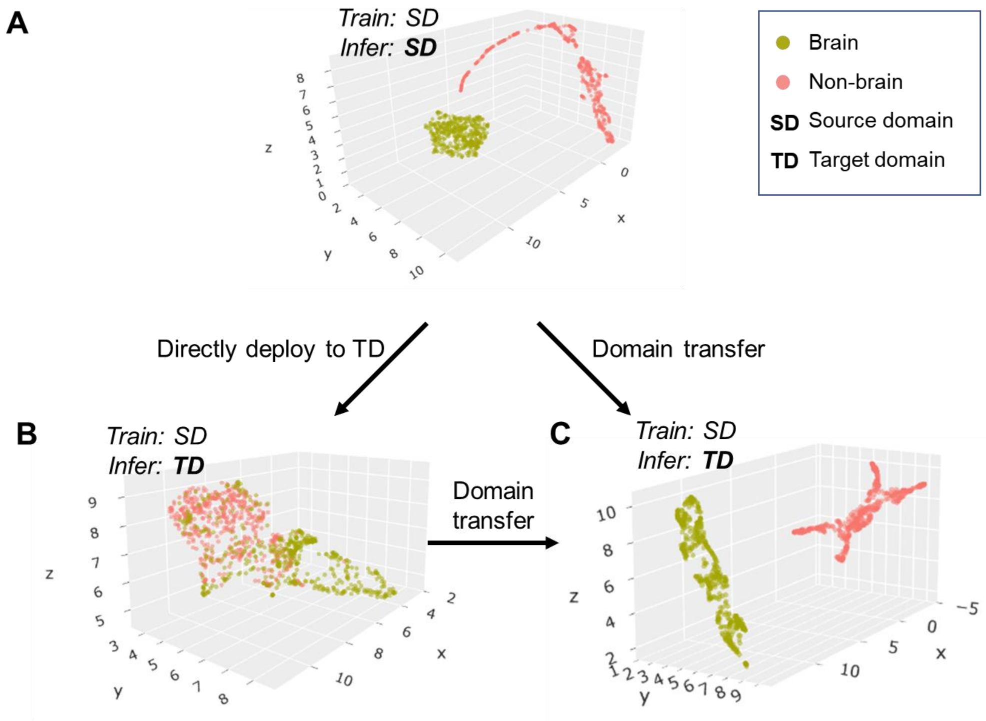


**Figure S11. UMAP visualizes BEN's transfer learning.**
**(A - C)** Scatter plot of neural network feature map clusters. An unsupervised clustering algorithm (UMAP) was used to visualize the semantic features. Different colors in the scatter plot indicate different clusters (green: brain, red: non-brain). Visualization of feature maps via UMAP during the following three intermediate processes: **(A)** semantic features of brain and non-brain are separate in the source domain; **(B)** these features are intermingled in the target domain without transfer learning; **(C)** features are separated again in the target domain after BEN's domain transferring strategy.

**Supplementary Tables**

**Table S1. MRI scan information of the fifteen animal datasets and three human datasets.**

FDU: Fudan University. UCAS: University of Chinese Academy of Sciences; UNC: University of North Carolina at Chapel Hill; ABCD: Adolescent Brain Cognitive Developmental study; ZIB: Zhangjiang International Brain BioBank at Fudan University.

| Species | Modality | Magnetic Field (T) | Scans | Slices | In-plane Resolution (mm) | Thickness (mm) | Manufacturer | Institution |
|---|---|---|---|---|---|---|---|---|
| Mouse | T2WI | 11.7 | 243 | 9,030 | 0.10*0.10 | 0.4 | Bruker | FDU |
| | | 9.4 | 14 | 448 | 0.06*0.06 | 0.4 | Bruker | UCAS |
| | | 7 | 14 | 126 | 0.08*0.08 | 0.8 | Bruker | UCAS |
| | EPI | 11.7 | 54 | 2,198 | 0.2*0.2 | 0.4 | Bruker | FDU |
| | | 9.4 | 20 | 360 | 0.15*0.15 | 0.5 | Bruker | UCAS |
| | SWI | 11.7 | 50 | 1,500 | 0.06*0.06 | 0.5 | Bruker | FDU |
| | ASL | 11.7 | 58 | 696 | 0.167*0.167 | 1 | Bruker | FDU |
| Rat | T2WI | 11.7 | 132 | 5,544 | 0.14*0.14 | 0.6 | Bruker | FDU |
| | | 9.4 | 55 | 660 | 0.1*0.1 | 1 | Bruker | UNC † |
| | | 7 | 88 | 4,400 | 0.09*0.09 | 0.4 | Bruker | UCAS |
| | EPI | 9.4 | 55 | 660 | 0.32*0.32 | 1 | Bruker | UNC † |
| **Sum of rodent** | | | **783** | **25,622** | | | | |
| Marmoset | T2WI | 9.4 | 62 | 2,480 | 0.2*0.2 | 1 | Bruker | UCAS |
| | EPI | 9.4 | 50 | 1,580 | 0.5*0.5 | 1 | Bruker | UCAS |
| Macaque * | T1WI | 4.7<br>3<br>1.5 | 76 | 20,063 | 0.3*0.3 ~0.6*0.6 | 0.3~0.75 | Siemens, Bruker, Philips | Multicenter |
| | EPI | 1.5 | 58 | 2,557 | 0.7*0.7~2.0*2.0 | 1.0~3.0 | Siemens, Bruker, Philips | Multicenter |
| **Sum of nonhuman primate** | | | **246** | **26,680** | | | | |
| Human (ABCD) | T1WI | 3 | 3,250 | 552,500 | 1.0*1.0 | 1 | GE<br>Siemens<br>Philips | Multicenter |
| Human (UK Biobank) | T1WI | 3 | 963 | 196,793 | 1.0*1.0 | 1 | Siemens | Multicenter |
| Human (ZIB) | T1WI | 3 | 388 | 124,160 | 0.8*0.8 | 0.8 | Siemens | FDU |
| Sum of human | | | 4,601 | 873,453 | | | | |
| **In total** | | | **5,630** | **925,755** | | | | |

* https://fcon_1000.projects.nitrc.org/indi/indiPRIME.html

† https://openneuro.org/datasets/ds002870/versions/1.0.0

**Table S2. Performance comparison of BEN with SOTA methods on the source domain (Mouse-T2WI-11.7T).**

Dice: Dice score; SEN: sensitivity; SPE: specificity; ASD: Average Surface Distance; HD95: the 95-th percentile of Hausdorff distance.

| Method | Dice | SEN | SPE | ASD | HD95 |
|---|---|---|---|---|---|
| Sherm | 0.9605 | 0.9391 | **0.9982** | 0.6748 | 0.4040 |
| AFNI | 0.9093 | 0.9162 | 0.9894 | 1.9346 | 0.9674 |
| FSL | 0.3948 | **1.0000** | 0.6704 | 20.4724 | 5.5975 |
| BEN | **0.9859** | 0.9889 | **0.9982** | **0.3260** | **0.1436** |

**Table S3. Performance comparison of BEN with SOTA methods on two public datasets.**
Dice: Dice score; Jaccard: Jaccard Similarity; SEN: sensitivity; HD: Hausdorff distance.

CARMI dataset * (Rodent)

T2WI

| Methods | Dice | Jaccard | SEN | HD (voxels) |
|---|---|---|---|---|
| RATS (Oguz et al., 2014) | 0.91 | 0.83 | 0.85 | 8.76 |
| PCNN (Chou et al., 2011) | 0.89 | 0.80 | 0.90 | 7.00 |
| SHERM (Liu et al., 2020) | 0.88 | 0.79 | 0.86 | 6.72 |
| U-Net (Hsu et al., 2020) | 0.97 | 0.94 | 0.96 | 4.27 |
| BEN | **0.98** | **0.95** | **0.98** | **2.72** |

EPI

| Methods | Dice | Jaccard | SEN | HD (voxels) |
|---|---|---|---|---|
| RATS (Oguz et al., 2014) | 0.86 | 0.75 | 0.75 | 7.68 |
| PCNN (Chou et al., 2011) | 0.85 | 0.74 | 0.93 | 8.25 |
| SHERM (Liu et al., 2020) | 0.80 | 0.67 | 0.78 | 7.14 |
| U-Net (Hsu et al., 2020) | 0.96 | 0.93 | 0.96 | 4.60 |
| BEN | **0.97** | **0.94** | **0.98** | **4.20** |

PRIME-DE † (Macaque)

T1WI

| Methods | Dice | Jaccard | SEN | HD (voxels) |
|---|---|---|---|---|
| FSL | 0.81 | 0.71 | 0.96 | 32.38 |
| FreeSurfer | 0.56 | 0.39 | **0.99** | 42.18 |
| AFNI | 0.86 | 0.79 | 0.82 | 25.46 |
| U-Net (Wang et al., 2021) | 0.98 | - | - | - |
| BEN | **0.98** | 0.94 | 0.98 | **13.21** |

* https://openneuro.org/datasets/ds002870/versions/1.0.0
† https://fcon_1000.projects.nitrc.org/indi/indiPRIME.html

**Table S4. Ablation study of each module of BEN in the source domain.**
(a) training with all labeled data using U-Net. The backbone of BEN is non-local U-Net (NL-U-Net). (b) training with 5% labeled data. (c) training with 5% labeled data using BEN's semi-supervised learning module (SSL). The remaining 95% of the unlabeled data is also used for the training. Since this ablation study is performed on the source domain, the adaptive batch normalization (AdaBN) module is not used. Dice: Dice score; SEN: sensitivity; SPE: specificity; HD95: the 95-th percentile of Hausdorff distance.

| Method | Scans used | | Metrics | | | |
|---|---|---|---|---|---|---|
| | Labeled | Unlabeled | Dice | SEN | SPE | HD95 |
| [a]U-Net | 243 | 0 | 0.9773 | 0.9696 | **0.9984** | 0.2132 |
| [a]Backbone | 243 | 0 | **0.9844** | **0.9830** | **0.9984** | **0.0958** |
| [b]U-Net | 12 | 0 | 0.9588 | 0.9546 | 0.9945 | 1.1388 |
| [b]Backbone | 12 | 0 | 0.9614 | 0.9679 | **0.9970** | 0.7468 |
| [c]Backbone + SSL | 12 | 231 | **0.9728** | **0.9875** | 0.9952 | **0.2937** |

**Table S5**. **Ablation study of each module of BEN in the target domain**.
(a) training from scratch with all labeled data. The backbone of BEN is non-local U-Net (NL-U-Net). (b) training from scratch with 5% labeled data. (c) fine-tuning (using pretrained weights) with 5% labeled data. (d) fine-tuning with 5% labeled data using BEN's SSL and AdaBN modules. The remaining 95% of the unlabeled data is also used for the training stage. Dice: Dice score; SEN: sensitivity; SPE: specificity; HD95: the 95-th percentile of Hausdorff distance.

| Method | Pretrained | Scans used | | Metrics | | | |
|---|---|---|---|---|---|---|---|
| | | Labeled | Unlabeled | Dice | SEN | SPE | HD95 |
| [a]Backbone (from scratch) | | 132 | 0 | **0.9827** | 0.9841 | 0.9987 | **0.1881** |
| [b]Backbone (from scratch) | | 7 | 0 | 0.8990 | 0.8654 | 0.9960 | 4.6241 |
| [c]Backbone | ✓ | 7 | 0 | 0.9483 | 0.9063 | **0.9997** | 0.6563 |
| [c]Backbone + AdaBN | ✓ | 7 | 0 | 0.9728 | **0.9875** | 0.9952 | 0.2937 |
| [d]Backbone + SSL | ✓ | 7 | 125 | 0.9614 | 0.9679 | 0.9970 | 0.7468 |
| [d]Backbone + AdaBN + SSL | ✓ | 7 | 125 | **0.9779** | 0.9763 | 0.9986 | **0.2912** |

**Table S6**. **BEN provides interfaces for the following conventional neuroimaging software**.

| Name | Link |
|---|---|
| AFNI | afni.nimh.nih.gov/afni |
| ANTs | stnava.github.io/ANTs/ |
| FSL | fsl.fmrib.ox.ac.uk/fsl/fslwiki |
| FreeSurfer | freesurfer.net |
| SPM | www.fil.ion.ucl.ac.uk/spm |
| Nipype | pypi.org/project/nipype/ |

**Table S7**. **Protocols and parameters used for conventional neuroimaging toolboxes**.

| Method | Command | Parameter | Description | Range | Chosen value |
|---|---|---|---|---|---|
| AFNI | 3dSkullStrip | -marmoset | Brain of a marmoset | on/off | on for marmoset |
| | | -rat | Brain of a rat | on/off | on for rodent |
| | | -monkey | Brain of a monkey | on/off | on for macaque |
| FreeSurfer | mri_watershed | -T1 | Specify T1 input volume | on/off | on |
| | | -r | Specify the radius of the brain (in voxel unit) | positive number | 60 |
| | | -less | Shrink the surface | on/off | off |
| | | -more | Expand the surface | on/off | off |
| FSL | bet2 | -f | Fractional intensity threshold | 0.1~0.9 | 0.5 |
| | | -m | Generate binary brain mask | on/off | on |
| | | -n | Don't generate the default brain image output | on/off | on |
| Sherm | sherm | -animal | Species of the task | 'rat' or 'mouse' | according to the task |
| | | -isotropic | Characteristics of voxels | 0/1 | 0 |
